# CRISPRi Screen Uncovers lncRNA Regulators of Human Monocyte Growth

**DOI:** 10.1101/2024.11.25.624758

**Authors:** Cristina Flores-Arena, Eric Malekos, Sergio Covarrubias, Lisa Sudek, Christy Montano, Valory Dempsey, Vuky Hyunh, Sol Katzman, Susan Carpenter

## Abstract

Long noncoding RNAs are emerging as critical regulators of biological processes. While there are over 20,000 lncRNAs annotated in the human genome we do not know the function for the majority. Here we performed a high-throughput CRISPRi screen to identify those lncRNAs that are important in viability in human monocytes using the cell line THP1. We identified a total of 38 hits from the screen and validated and characterized two of the top intergenic hits. The first is a lncRNA neighboring the macrophage viability transcription factor IRF8 (*RP11-542M13*.*2* hereafter referred to as long noncoding RNA regulator of monocyte proliferation, *LNCRMP*) and the second is a lncRNA called *OLMALINC* (oligodendrocyte maturation-associated long intervening non-coding RNA) that was previously found to be important in oligodendrocyte maturation. Transcriptional repression of *LNCRMP and OLMALINC* from monocytes severely limited their proliferation capabilities. RNA-seq analysis of knockdown lines showed that *LNCRMP* regulated proapoptotic pathways while knockdown of *OLMALINC* impacted genes associated with the cell cycle. Data supports both *LNCRMP* and *OLMALINC* functioning *in cis* to regulate their neighboring proteins that are also essential for THP1 cell growth. This research highlights the importance of high-throughput screening as a powerful tool for quickly discovering functional long non-coding RNAs (lncRNAs) that play a vital role in regulating monocyte viability.

## Introduction

Monocytes are the largest white blood cell circulating in the periphery that play critical roles in regulating innate immune responses (1). Key features of monocytes are their ability to sense pathogen-associated molecular patterns, proliferate and migrate to tissues where they differentiate into macrophage or dendritic cells (2, 3). Both monocytes and macrophages are critical to the activation and resolution of inflammation through phagocytosis, initiation of cytokine and chemokine signaling cascades, and activation of the adaptive immune system (3, 4). Under normal homeostatic conditions, circulating monocytes are short lived and die by apoptosis (4). Increased accumulation of monocytes can promote diseases including cancer, cardiovascular disease, liver fibrosis and multiple sclerosis liver (5). Considering the vast number of biological processes monocytes take part in, including development, wound healing, cancer progression (4, 6) we sought to determine what role long noncoding RNAs might play in monocyte growth and proliferation.

Long noncoding RNAs represent the largest group of RNA produced from the human genome. LncRNAs are defined as transcripts longer than 500 nucleotides, often spliced and polyadenylated, but without the ability to code for proteins (7). In recent years, research has revealed diverse roles for lncRNAs in various biological processes, such as cell differentiation, cancer, and immunity (8–11). However, the functions of the vast majority of lncRNAs remain unknown. A significant hurdle in studying lncRNAs has been the lack of reliable and specific methods for targeting them, especially in high-throughput settings. Traditional CRISPR/Cas9 techniques, which rely on inducing frameshift mutations, are ineffective against lncRNAs as they lack open reading frames. The emergence of CRISPR inhibition (CRISPRi), a modified CRISPR/Cas9 technology, has provided a powerful solution. CRISPRi can silence lncRNA transcription by inducing repressive chromatin marks at their start sites, making it a valuable tool for uncovering lncRNA functions. While lncRNA expression profiles have been examined in human monocytes (12) there is a lack of functional data and a small number of lncRNAs have been identified as playing roles in monocyte differentiation, there is a lack of systematic approaches to identify those lncRNAs important in monocyte survival (13, 14)

Liu, Horlbeck et al published one of the first lncRNA screens aimed at understanding which lncRNAs are essential in human cells (15). They profiled 7 cell lines identifying a total of 499 lncRNA using high throughput pooled CRISPRi screening. Interestingly the majority of the hits were unique to each cell type indicating that lncRNAs are highly cell and tissue specific in expression and function. We recently utilized a similar approach to identify all lncRNAs involved in monocyte induced inflammation and monocyte to macrophage differentiation (9, 16). Here we applied a similar approach utilizing our THP1 specific lncRNA library (16) to profile and determine which lncRNAs are important to monocyte survival and proliferation. Of the top 35 hits involved in cell monocyte cell viability, 18 are antisense transcripts, 8 are intergenic, 4 are now annotated as protein coding, 3 are intronic and 2 are bidirectional transcripts. We perform mechanistic studies on two hits, *LNCRMP* and *OLMALINC*. RNA-sequencing following knockdown of each lncRNA reveals that they both function to regulate viability in different ways. While both genes can impact their neighbor *in cis OLMALINC* regulates cell cycle pathways while *LNCRMP* impacts interferon signaling as well as pro-apoptotic proteins. Here we present the power of CRISPR screening to identify functional lncRNAs important in cell growth in human monocytic cells.

## Results

### High Throughput CRISPR screening identifies lncRNAs involved in THP1 monocyte cell growth

In order to identify lncRNAs important in the regulation of monocytic cell growth, we took our recently generated monocytic specific lncRNA CRISPR library and infected THP1 cells (16). The library contains ∼25,000 individual single guide RNAs (sgRNAs) with 10 sgRNAs targeting the transcription start sites of 2,342 lncRNAs annotated in the human genome assembly GRCh37 (hg19). The same design and cloning strategy were used as previously described (15) and is presented in Figure 1A. THP1s were infected and selected with puromycin for 7 days. After 21 days, genomic DNA was extracted and compared to the starting library to identify the sgRNAs that dropped out or enriched over time. Hits were ranked by comparing fold-change of TSS-targeting sgRNAs to non-targeting control sgRNAs and assigning each a z-score (Fig.1B). Genes with a score of less than -3 were defined as significant positive regulators of THP1 growth while genes above 3 were defined as significant negative regulators of growth (Fig. 1B). Highlighted in the figure are positive control gene hits *PVT1* and *CASP8AP2. PVT1* was previously shown to act as a growth suppressor and *CASP8AP2* is a viability gene (17–19). Here we are only focusing on the sgRNAs that dropped out representing positive regulators of growth in THP1 cells. In Figure 1.C we rank the top 35 hits based on the three performing sgRNAs (Fig.1C, Table 1). 18 lncRNA hits are antisense to a protein coding gene, 8 hits are intergenic, defined as having their own promoters at least 1kb away from promoters of neighboring genes, 4 hits are protein coding (previously annotated as noncoding in older genome build), 3 are intronic and 2 are from a bidirectional promoter (Fig.1D). One of the top hits was CASP8AP2 which was previously identified in a CRISPRi screen as an essential viability factor in lung cancer (Fig.1E) (18). For the remainder of this study, we focus on two hits which were selected as they are both intergenic long noncoding RNAs that have not been implicated in monocyte or macrophage biology previously (Fig.1E). *OLMALINC* neighbors the protein coding gene stearoyl-CoA desaturase (SCD) and has previously been shown to regulate SCD *in cis* in neuronal cells (20). Interestingly SCD itself has previously been reported to be a viability gene in THP1s in the genome CRISPR screens database (Fig.1E) as well as in the Broads depmap portal https://depmap.org/portal/cell_line/ACH-000146 (gene effect in THP1s of -1.081222 and a z score of -0.63). The second lncRNA RP11-542M13.2 (AC092723.1), which we call lncRNA regulating monocyte proliferation - *LNCRMP* - was selected as it neighbors the important protein coding gene IRF8. We previously reported that IRF8 is a macrophage viability gene in mouse bone marrow derived macrophages, and indeed it has also been reported to act as a viability gene in THP1 screens(21) and Fig1E and in the depmap portal with a gene effect in THP1s of -1.18 and a Z score of -7.85.

**Figure 1:**
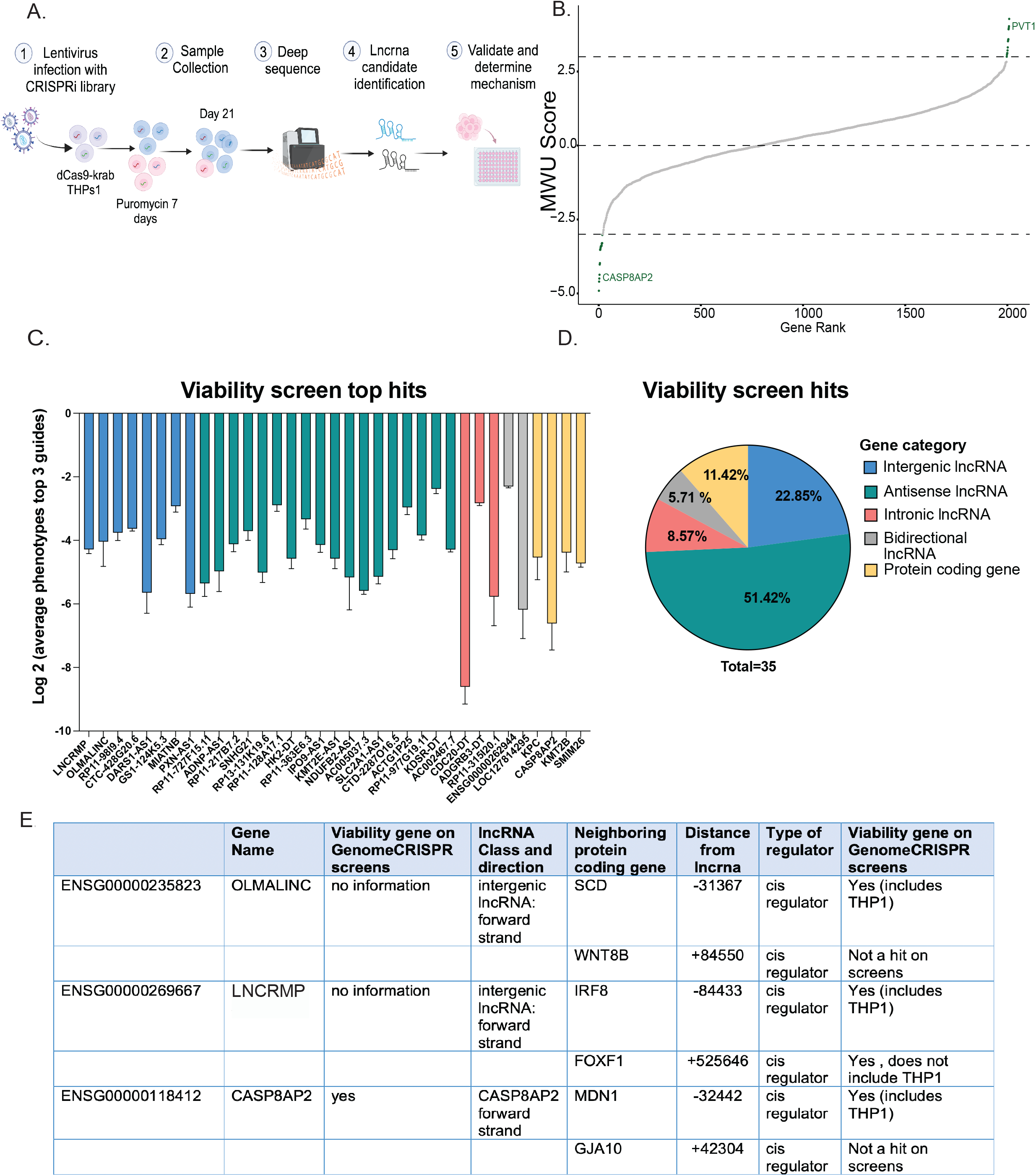
CRISPRi screen identifies lncRNAs that impact macrophage cell growth (A) Schematic of drop out CRISPRi screen pipeline. sgRNAs were designed to target the transcription start sites of over 2000 Gencode hg19 annotated lncRNAs. Transcription start sites were predicted using data from FANTOM and ENCODE. THP1 lncRNA expression was estimated from THP1 RNAseq data. NFkB-EGFP-CRISPRi-THP1 cells were infected with pooled sgRNA libraries.The cells were selected with puromycin and samples were collected at day 21 to identify the sgrnas that dropped out from the original library. sgRNAs from the library and day 21 were PCR amplified and sequenced **(B) Screen hits**. Mann Whitney U analysis was used to determine the gene ranking from the RNA seq data. **(C) Top performing sgRNAs**. Top 3 guides for each significant hit are represented and color coded to match their gene category in D. **(D) LncRNA hit categories**. Pie chart for the different categories of lncrnas in the CRISPRi screen. Intergenic lncrnas are lncrnas localized between two genes and they have their own transcription start site (TSS). Antisense lncrnas are lncrnas that transcribed from the same gene locus in the opposite direction. Intronic lncrnas are localized within an intron of a gene. Bidirectional lncrnas are 1kb away from the gene locus and they transcribed in the opposite direction. **(E) Table** summarizing top candidate lncRNAs, *OLMALINC* and *LNCRMP* and a positive control hit *CASP8AP2*.

### Validation of the top hits *OLMALINC* and *LNCRMP* as regulators of THP1 monocyte growth

We sought to better characterize *OLMALINC* and *LNCRMP*, our candidate hits. To this end we generated three independent sgRNA lines targeting each gene and confirmed knockdown by qPCR compared to a scrambled control line (Fig. 2A-B) consistently achieving >95% knockdown of *OLMALINC*, and 50-60% for *LNCRMP*. Due to these genes being associated with cell growth and viability, all phenotypes were assessed after 10 days of puromycin selection to minimize selection of cells without knockdown. After validating CRISPRi knockdown we performed a mix cell assay to validate these genes as regulators of cell growth (Fig 2C). We combined cherry positive cells (containing sgRNAs) with cherry negative cells at a 1:1 ratio and monitored cell growth over time as assessed by changes in the ratio of cherry positive to cherry negative cells as measured by flow cytometry. All three sgRNAs targeting either *OLMALINC, LNCRMP* or our positive control *CASP8AP2*, resulted in between 75-95% decreased cell growth compared to the 5 negative control sgRNAs (Fig 2D). This sensitive assay confirmed that both *OLMALINC* and *LNCRMP* contribute to THP1 cell growth.

**Figure 2:**
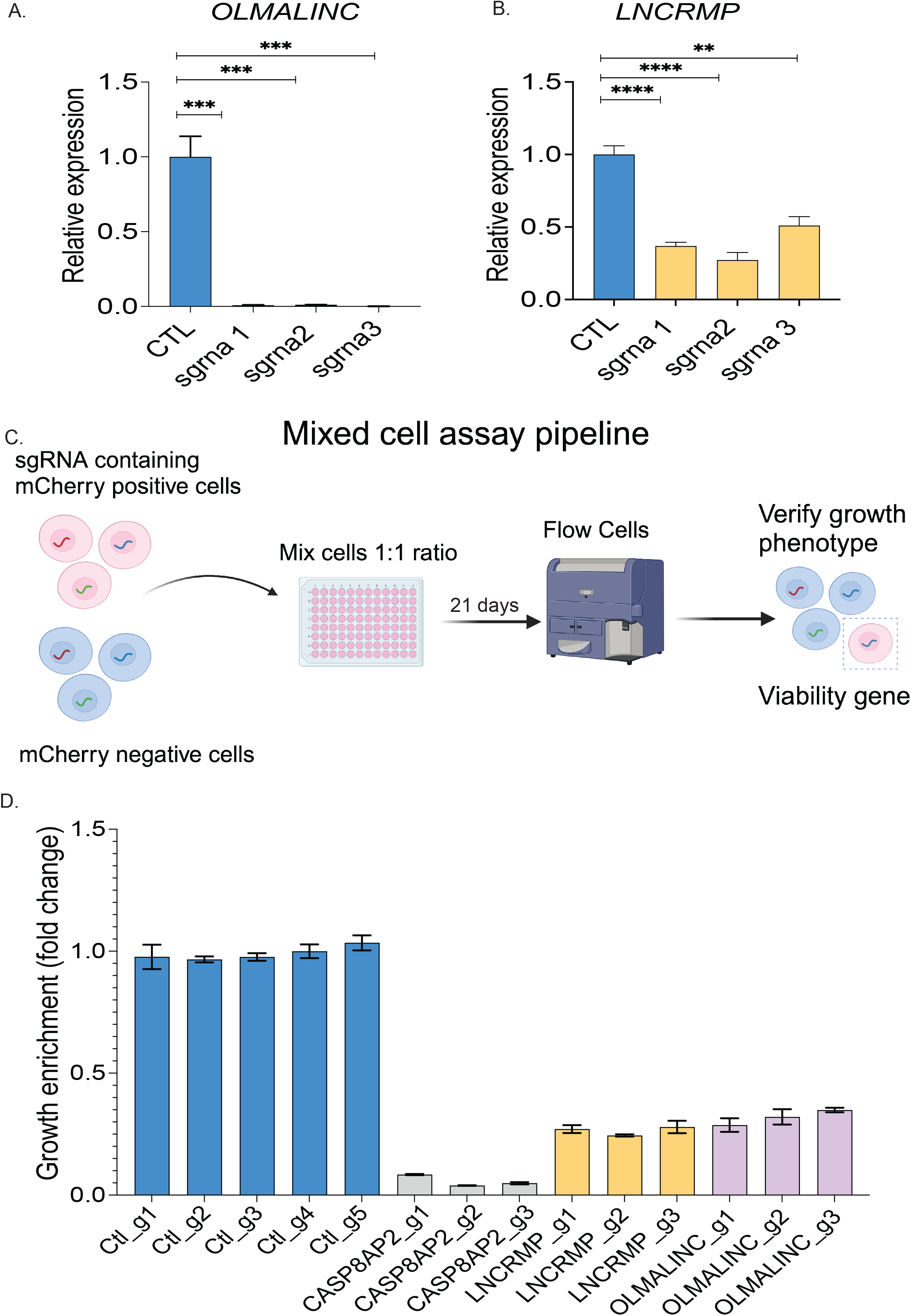
Validation of *OLMALINC* and *LNCRMP* as hits involved in macrophage growth CRISPRi knockdown of *OLMALINC* and *LNCRMP* (A-B). sgRNAs (sgRNAs 1, 2, 3) were designed to target top lncRNA candidates *OLMALINC* and *LNCRMP*. qPCR measurement of both genes across three replicate experiments shows knockdown of *OLMALINC (A) and LNCRMP (B)* by all three sgRNAs (p-values<0.05) vs a non-targeting control sgRNA **(C) Schematic of macrophage growth validation assay**. Mcherry positive cells sgRNA containing cells were mixed with mcherry negative cells in a 1:1 ratio and plated in triplicates for each sgrna for each gene. The cells were flowed every other day for 21 days. Mcherry enrichment was then calculated to determine the cell growth. **(D) Mixed cell assay results for *OLMALINC* and *LNCRMP***. We combined cherry positive cells (containing sgRNAs) with unedited cherry negative cells at a 1:1 ratio and monitored growth of sgRNA infected cells (cherry) relative to uninfected reference cells in a mix-cell growth assay for 21 days. Experiment was repeated 3 times and a representative experiment was displayed.

### *OLMALINC* is a cytosolic gene regulating monocyte growth via SCD and Immune/Cell cycle checkpoints

*OLMALINC* has previously been reported to act *in cis* regulating its neighboring coding gene, SCD (20, 22). To determine if there could be a similar mechanism at play in our THP1 cell system we first performed RNA-sequencing comparing negative control sgRNAs to three sgRNAs targeting *OLMALINC* outlined in the UCSC browser track in Fig.3A. We saw robust knockdown of *OLMALINC*, consistent with our qPCR results (Fig.3A-B). Expression levels of *OLMALINC*’s neighboring protein coding gene, *SCD*, were also significantly reduced, consistent with *cis* regulatory activity. *Cis* acting elements are known to frequently occupy the same TADs as their target, therefore we looked at THP1 HiC data and confirmed that SCD and *OLMALINC* share a TAD as determined by HiC interaction data and consistent with TAD boundaries determined by CTCF peaks (Fig. 3D). This is consistent with a previous study indicating that *OLMALINC* works *in cis* through an enhancer RNA mechanism (22). Knockdown of *OLMALINC* resulted in dysregulation of a number of protein coding genes (Fig. 3E), including proteins such as DBF4-CDC7 kinase regulatory subunit A (DBF4) and CCN family member 4,CCNA4 involved in cell cycle control (Fig. 3F) and Interferon Induced proteins with tetratricopeptide repeats (IFITs) and 2′,5′-oligoadenylate synthetase (OAS) proteins involved in Interferon signaling (Fig. 3G).

**Figure 3:**
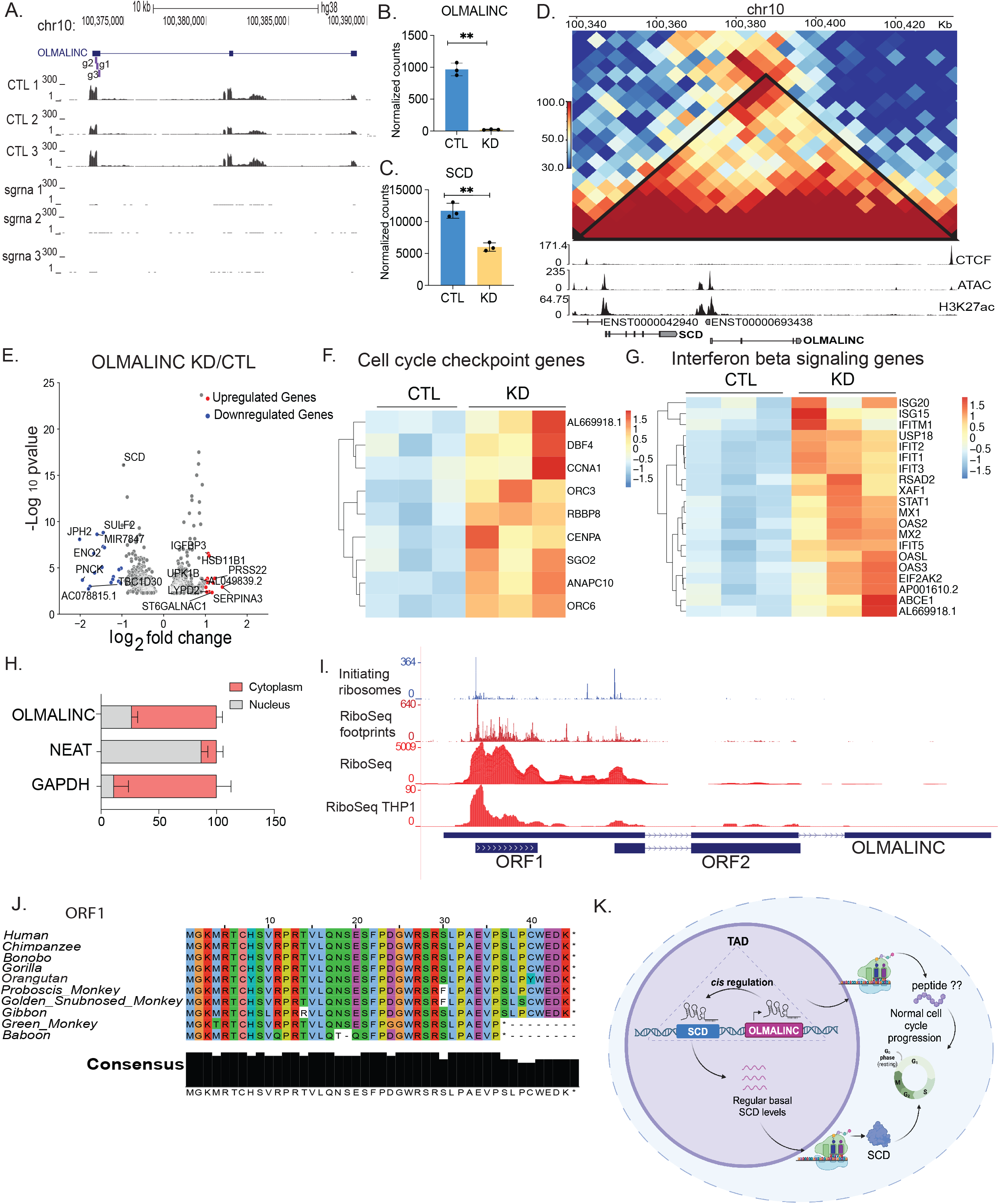
Functional characterization of *OLMALINC’s* regulation of monocyte growth. (A-C). **CRISPRi knockdown of *OLMALINC* impacts SCD expression** (**A**) UCSC genome browser track displays RNA-Seq reads from control (Ctl) and *OLMALINC* knockdown (KD) cell lines at the *OLMALINC* genomic locus. (**B-C**) Normalized read counts from DEseq analysis for *OLMALINC* (B) and its neighboring protein SCD (C). **(D) *OLMALINC* and *SCD* share a TAD**. *OLMALINC* and SCD are part of the same topological associated domain (TAD). Indicating a close genomic proximity and cis regulation. **(E) Differentially expressed genes following *OLMALINC* knockdown**. Volcano plot displaying significantly upregulated and downregulated genes in *OLMALINC* KD cells. (**F-G**) Heatmaps of top 20 upregulated or downregulated genes in *OMALINC* KD cells and control cells. **(H) *OLMALINC* is expressed in the cytosol**. RT-qPCR analysis of *OLMALINC* expression in cytoplasmic and nuclear fractions from THP1 cells.**(I) *OLMALINC* possibly encodes short peptides**. Riboseq data for *OLMALINC* shows Riboseq aggregates at open reading frame 1 (ORF1) and conservation between different species. **Track 1** shows initiating Riboseq aggregates in the different ORFs. **Track 2** shows Riboseq footprint aggregates. **Track 3** shows Riboseq coverage aggregate. **Track 4** shows Riboseq coverage in THP1 cells. **(J) *OLMALINC* Peptide conservation**. Consensus conserved sequence is observed in different species such as human, gorilla and green monkey. **(K) Proposed working model** Briefly, *OLMALINC* acts as a *cis* regulator of SCD (viability coding gene) and regulates its expression. Under basal conditions, *OLMALINC* is expressed at basal levels and regular levels of SCD are expressed which results in a normal cell cycle. In the absence of *OLMALINC*, SCD levels are lower and cell cycle is altered impacting cell growth.

To gain further functional insight into *OLMALINC* we first determined its localization by fractionation of THP1 cells followed by qRT-PCR. NEAT1 was measured as a nuclear control and GAPDH served as the cytosolic control (Fig. 3H). *OLMALINC* is dominantly expressed in the cytosol with some expression (∼30%) in the nuclear compartment (Fig. 3H). Considering the high levels of *OLMALINC* localized in the cytosol we next asked if the locus is indeed noncoding or possibly encoding a short encoded peptide. Previous work by Presener et al, identified *OLMALINC* as a gene important in viability in cancer cells. They also showed that it harbored an open reading frame (ORF) and active Cas9 targeting the coding region affected viability in HELA cells (23). Indeed Riboseq, including initiating ribosome reads, Riboseq footprints aggregated by GWIPS, and Riboseq from THP1 cells indicated that there are two translated ORFs within the *OLMALINC* locus. In THP1s, the strongest Riboseq signals support open reading frame 1 (ORF1) and we find that this peptide sequence is highly conserved across primates (Fig. 3J). This data suggests that there could be dual roles for this gene locus. Nuclear *OLMALINC* could function *in cis* to regulate its neighboring protein coding gene. It is also possible that the functional peptides contribute THP1 cell growth either through regulating SCD or other gene networks (Fig. 3K).

### *LNCRMP* is expressed in both the cytosol and nucleus and regulates both its neighboring gene IRF8 and other IFN genes

To better understand *LNCRMP*’s role in regulating monocyte growth we performed RNA-seq to determine what genes are altered when *LNCRMP*s expression is reduced. UCSC Browser tracks of the *LNCRMP* locus confirmed successful knockdown in the two sgRNAs utilized compared to negative controls (Fig. 4A). Normalized read counts from RNA-seq indicate successful knockdown of *LNCRMP* which resulted in small but significant downregulation of its neighboring gene *IRF8*. Utilizing HiC data we found that both *LNCRMP* and *IRF8* occupy the same TAD within THP1s and is suggestive of a *cis* regulatory effect between the two genes. Figure 1 D shows a volcano plot of dysregulated genes following *LNCRMP* knockdown. Significantly upregulated genes with a log_2_ fold change of 1 are in red and downregulated genes with a log_2_ fold change of -1 are in blue. The top 20 upregulated genes in the *LNCRMP* deficient cells fall into the defense and IFN signaling GO terms indicative of a role for *LNCRMP* in regulating this pathway. Using fractionation followed by qRT-PCR we found that *LNCRMP* is expressed in both the nuclear and cytosolic compartments. Given *LNCRMPs* expression in the cytosol we examined THP1 Riboseq data but did not identify any significant reads in the region suggestive of *LNCRMP* being a bonafide noncoding RNA (Supp. Fig.1). Previously we identified IRF8 as a viability gene in mouse macrophages and our proposed model based on the current findings is that *LNCRMP* can function *in cis* to regulate IRF8 which could have a downstream impact resulting in monocytic cell death. This is consistent with previous reports on this lncRNA that showed it can function to regulate its neighbor IRF8 through an enhancer RNA mechanism involving recruitment of TET1 to the IRF8 promoter regulating IRF8 expression by affecting methylation levels in the U937 monocytic cell line (24).

**Figure 4:**
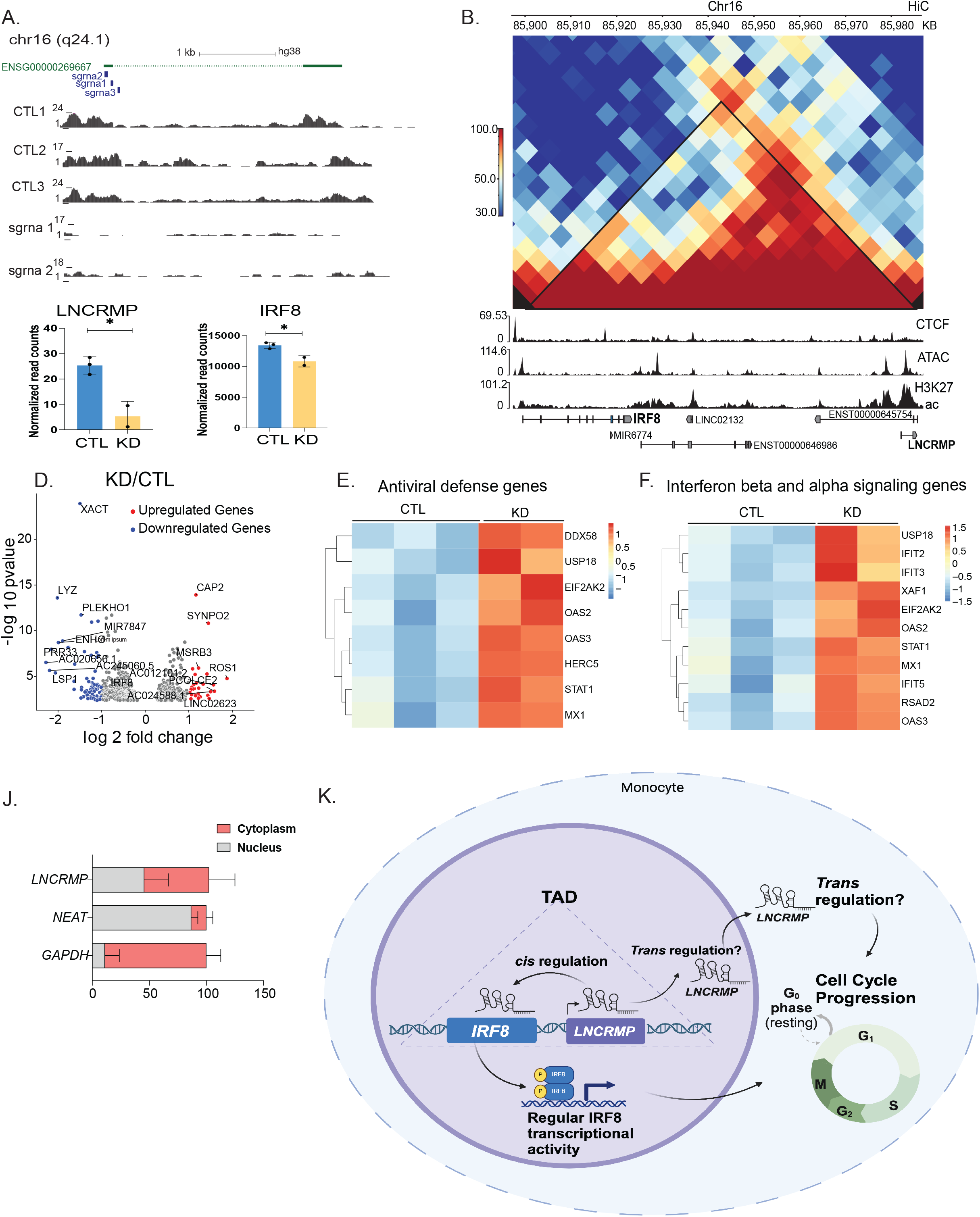
Characterization of *LNCRMP*’s role in monocytic growth and function. **(A)** UCSC genome browser track displays RNA-Seq reads from control (Ctl) non-targeting sgRNAs and *LNCRMP* knockdown (KD) cell lines at the *LNCRMP* genomic locus. **(B)** Expression of *LNCRMP* (top graph) and its neighboring protein SCD (bottom graph) from RNA-seq data comparing Ctl sgRNAS to *LNCRMP* sgRNA. **(C)** *LNCRMP* and IRF8 are part of the same topological associated domain (TAD). Indicating a close genomic proximity and possible cis regulation. **(D)** Volcano plot displaying significantly upregulated and downregulated genes in *LNCRMP* KD cells. **(E-F)** Heatmaps of top upregulated *LNCRMP* KD cells and controls across two categories outlined. **(G)** RT-qPCR analysis of *LNCRMP* expression in cytoplasmic and nuclear fractions from THP1 cells. **(H)** Proposed working model of how *LNCRMP* acts as a regulator of THP1 viability. Briefly, *LNCRMP* acts in *cis* to regulate IRF8

## Discussion

Here we present a CRISPRi-based screen in the human monocytic cell line THP1 identifying lncRNAs that regulate cell growth. We focused on two top hits: *LNCRMP* and *OLMALINC*. Interestingly, neither lncRNA was identified in a previous screen of 499 lncRNAs involved in cell growth across 7 different cell lines (15). This discrepancy likely reflects cell type-specific roles for these lncRNAs. *LNCRMP* was not expressed in any of the cell lines tested by Liu et al., while *OLMALINC* was expressed at varying levels (TPMs 0.7-29) but did not significantly affect growth in those contexts. These findings underscore the importance of cell type context in lncRNA function, consistent with Liu et al.’s observation that cell type-specific expression was a major determinant of growth regulation. For example, they found that *LINC00263*, despite widespread expression, was only required for growth in U87 cells (15). It is also possible that other cell types have built in redundancy and therefore are not reliant on a particular lncRNA.

We have previously reported that protein coding genes can play specific roles in controlling growth in macrophages (21). We performed a whole genome wide active CRISPR screen and identified a total of 61 macrophage specific protein hits (21). Interestingly one of the hits is the transcription factor IRF8. IRF8 is a key transcription factor that plays roles in macrophage development, can act as a tumor suppressor and function to regulate inflammatory signaling (25–28). The locus surrounding human IRF8 is highly complex containing multiple possible lncRNA genes according to Gencode annotations. Despite this, we only identified the region encoding *LNCRMP* as a hit in THP1s. It is possible that the other lncRNAs play functional roles in other cell types. Interestingly a previous report by Zhou et al., identified a Lupus disease associated variant rs2280381 which lies 300bp downstream of *LNCRMP (*AC092723.1). Through mechanistic studies they showed that this variant impacted IRF8 and *LNCRMP* expression through regulation of binding of the transcription factor PU.1. They went on to show that *LNCRMP* recruits TET1 to the IRF8 promoter and affects expression levels through DNA methylation. Our data supports *LNCRMP* functioning as a growth regulator of monocytes most likely through its *cis* regulation of IRF8. Previous work on conditional knockdown of IRF8 in BMDMs resulted in upregulation of many genes, again supporting the idea that most of the effects of knocking down *LNCRMP* are through cis regulation of IRF8 (29). The effect we observed in the *LNCRMP* knockdown cells on IRF8 is modest. However, we had difficulty obtaining cell populations containing over 60% knockdown of *LNCRMP* (Fig. 2), perhaps because of the lethal effects of knocking this gene down resulting in loss of these cells from our population. It is also possible that there is a different start site for this gene in monocytes than the one annotated in gencode. If this is the case the targeting of our sgRNAs might not be optimal and impact the levels of knockdown obtained. Given that *LNCRMP* is expressed in both the cytosol and the nucleus it is possible that it has effects *in trans* outside of its role in regulating IRF8 *in cis*. Further work will be needed to fully understand the complexities of the locus and the role that *LNCRMP* plays in regulating IRF8 in myeloid cells. This work could be important in particular as it relates to the role that these genes play in regulating disease pathogenesis in conditions such as Lupus.

*OLMALINC* appears to be more widely expressed across cell types but has only been reported to play a role in regulating cell growth in the brain and now in monocytic cells (20). *OLMALINC* could impact different signaling pathways in various cell types possibly through its locus being multifunctional. We find it dominantly expressed in the cytosol of monocytes and have identified putative small open reading frames which can be translated to produce a short encoded peptide (SEP). A previous study also found *OLMALINC* to be a growth regulator in HELA cells (23) and they tiled the length of the gene using active Cas9 and found the regions containing ORF2 affected the growth phenotype again providing more evidence for a functional SEP within the *OLMALINC* locus. In the THP1 Riboseq data we analyzed, a more upstream ORF shows strong evidence of translation, implicating *OLMALINC* as a bicistronic RNA (Fig 3I). It is possible that its RNA plays roles in regulating its neighboring gene, SCD, *in cis* as previously reported (20). It is also possible that it functions to regulate SCD and other genes through its SEP thereby impacting monocytic cell growth. Again, further mechanistic work is required to fully understand the regulatory functions of this locus across cell types.

A recent study used RNA-targeting Cas13 to pinpoint long non-coding RNAs (lncRNAs) crucial for cell survival and growth (30). They systematically targeted lncRNAs in five cell lines, including THP1 cells, and found 778 essential lncRNAs. One key finding was that most of these lncRNAs function independently of nearby protein-coding genes. This highlights a key difference between DNA-targeting Cas9 and RNA-targeting Cas13: Cas13 allows researchers to study lncRNAs that act *in trans* while we observed that CRISPRi targeting dominated for antisense transcripts and *cis* regulatory lncRNAs. *LNCRMP* was not included in their lncRNA analysis annotation pipeline however they did identify *OLMALINC* (annotated as Hum_XLOC_005888) as a viability hit in THP1 cells. They also showed it was a significant hit in Hap cells, k562 and MDS-MB-231s but not in 293 cells.

This work demonstrates the power of CRISPR screening for rapidly identifying functional lncRNAs, a crucial step towards understanding this largely uncharacterized class of genes. Our findings, along with other recent studies, highlight the importance of *cis*-regulation by lncRNAs in fine-tuning complex signaling pathways. Notably, a recent study identified the first case of a neurological genetic disease caused by the loss of a single copy of the lncRNA CHASERR (31). This haploinsufficiency resulted in significant upregulation of a neighboring protein coding gene, leading to disease. It is likely that many lncRNAs play similar roles in fine-tuning the expression of neighboring genes, with potential implications for understanding the pathogenesis of complex inflammatory diseases. Further investigation of these multifunctional lncRNA loci is critical for unlocking new therapeutic targets and advancing our understanding of human health.

## Methods

### Cell lines

Wildtype (WT) THP1 cells were obtained from ATCC. All THP1 cell lines were cultured in RPMI 1640 supplemented with 10% low-endotoxin fetal bovine serum (ThermoFisher), 1X penicillin/streptomycin, and incubated at 37°C in 5% CO2.

### Lentivirus production

All constructs (table S9) were cotransfected into HEK293T cells with lentiviral packaging vectors psPAX (Addgene cat#12260) and pMD2.g (Addgene cat#12259) using Lipofectamine 3000 (ThermoFisher cat# L3000001) according to the manufacturer’s protocol. Viral supernatant was harvested 72 h post transfection.

### THP1--dCasKRAB-sgRNA

CRISPRi-THP1 cells were lentivirally infected with sgRNAs. sgRNA constructs were made from a pSico lentiviral backbone driven by an EF1a promoter expressing T2A flanked genes: puromycin resistance and mCherry. sgRNAs were expressed from a mouse U6 promoter. 20-Nucleotide forward/reverse gRNA oligonucleotides were annealed and cloned via the AarI site.

### Screening protocol

#### sgRNA library design and cloning

10 sgRNAs were designed for each TSS of hg19 annotated lncRNAs expressed in THP1s at baseline and upon stimulation. The sgRNA library also included 700 non-targeting control sgRNAs, and sgRNAs targeting 50 protein coding genes as positive controls. The sgRNA library was designed and cloned as previously described in (16).

### Screen analysis

THP1-CRISPRI cells were infected at a low multiplicity of infection (0.3) and selected with puromycin (10mg/ml) for 7 days. Cells were grown for a total of 21 days and maintained at at least 1000X coverage throughout. Genomic DNA was extracted and libraries prepared and sequenced as previously described (16). fastq.gz files were analyzed using the gRNA_tool: https://github.com/quasiben/gRNA_Tool. All guide RNA (sgRNA) + barcode reads were collapsed to obtain raw sgRNA counts. Counts were normalized to the median and fold-changes were calculated for each sgRNA. To identify significant genes for the growth screen, the

Mann-Whitney U test was performed comparing fold-changes for sgRNAs targeting each gene to non-targeting controls (described in (32)). For the growth screen, the Day 21 sample was compared to the library plasmid pool. Data is available at GSE278486

### Sequencing Data

RNA sequencing was performed to compare negative control THP1 cells to those with *OLMALINC* (labeled KO 823) or *LNCRMP* (labeled KO 667) knockdown. Data is available at GSE278485.

### Mixed cell growth assay

We used the top three performing sgrna guides from the screen to validate the lncrna genes. Briefly, sgRNA-infected THP1 cells (cherry-pos) were mixed with uninfected THP1 cells (cherry-neg) at a 1:1 ratio in triplicate. We used Flow cytometry to monitor the ratio of cherry-pos to cherry-neg cells at 0- and 21-days post plating. All validation cytometry was performed on the Attune NxT Flow Cytometer.

### Nuc/Cyt fractionation and RT-qPCR

WT THP1 cells were fractionated according to the NE-PER kit protocol (ThermoFisher cat# 78833) with RNAse inhibitor (Superase-IN, ThermoFisher cat# AM2696) added to the cytosolic and nuclear lysis buffers. 3 volumes of Trizol (TRI Reagent, Sigma T9424) were added to the fractions and RNA was isolated using the DIrect-zol RNA Miniprep Plus Kit (Zymo cat# R2052). 16uL of RNA isolated from fractions was reverse transcribed (iScript cDNA synthesis kit, Bio-Rad cat# 1708840) followed by qPCR (iTaq SYBRgreen Supermix, Bio-Rad cat# 1725121) using the cycling conditions as follows: 50C for 2 min, 95C for 2 min followed by 40 cycles of 95C for 15s, 60C for 30s and 72C for 45s.

**Supplemental Figure 1: *LNCRMP* does not encode any short peptides. (A**,**B)** RNASeq and Riboseq data for *IRF8* and *LNCRMP. LNCRMP* does not show Riboseq aggregates indicating there is no translation of this gene.

## Supporting information

SuppFig1

## Acknowledgments

C. F. A. was supported by [R35GM137801] and [K12GM139185] from the NIH,. C. M. is funded by a supplemental award to [R35GM137801] from NIGMS.. E.M. is supported by F31AI179201 from NIAID. S. C is supported by R35GM137801 from NIGMS

We would like to thank Dr. Alice Devigne of the UCSC CRISPR Core RRID:SCR_021207 for all her help with cell line generation.

