## Supplementary figures and images for "CRISPRi Screen Uncovers lncRNA Regulators of Human Monocyte Growth"

### SuppFig1

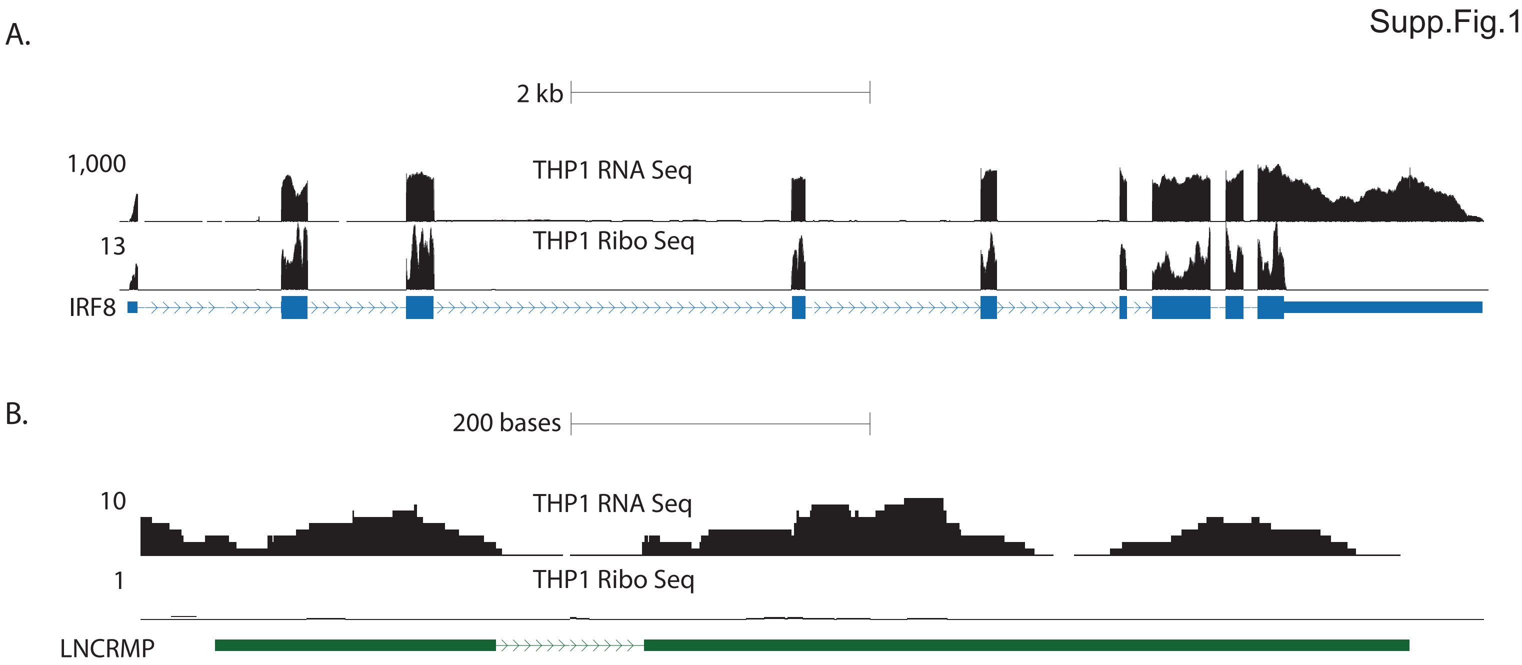
